# Placental nanoparticle gene therapy normalizes gene expression changes in the fetal liver associated with fetal growth restriction in a fetal sex-specific manner

**DOI:** 10.1101/2022.09.26.509494

**Authors:** Rebecca L Wilson, Kendal K Stephens, Helen N Jones

**Author notes:** **Corresponding Author:** Rebecca Wilson, Center for Research in Perinatal Outcomes, University of Florida, Gainesville, Florida 32610.

## Abstract

Fetal growth restriction (FGR) is associated with increased risk of developing Non-Communicable Diseases. We have a placenta-specific nanoparticle gene therapy protocol that increases placental expression of *human insulin-like growth factor 1* (*hIGF-1*), for the treatment of FGR *in utero*. We aimed to characterize the effects of FGR on hepatic gluconeogenesis pathways during early stages of FGR establishment, and determine whether treatment of the placenta with nanoparticle mediated *hIGF-1* therapy could resolve differences in the FGR fetus. Female Hartley guinea pigs (dams) were fed either a control or maternal nutrient restriction (MNR) diet using established protocols. At GD30-33, dams underwent ultrasound guided, transcutaneous, intra-placental injection of *hIGF-1* nanoparticle or PBS (sham), and were sacrificed 5 days post-injection. Fetal liver tissue was fixed and snap frozen for morphology and gene expression analysis. In female and male fetuses, liver weight as a percentage of body weight was reduced by MNR, and not changed with *hIGF-1* nanoparticle treatment. In female fetal livers, expression of *hypoxia inducible factor 1* (*Hif1α*) and *tumor necrosis factor* (*Tnfα*) were increased in MNR compared to Control, but reduced towards Control in MNR + *hIGF-1* livers. In male fetal liver, MNR increased expression of *Igf-1*, and decreased expression of *Igf-2* compared to Control. *Igf-1* and *Igf-2* expression was restored to Control levels in the MNR + *hIGF-1* group. This data provides further insight into the sex-specific mechanistic adaptations seen in FGR fetuses, and demonstrates that disruption to fetal developmental mechanisms may be returned to normal by treatment of the placenta.

## Introduction

Fetal growth restriction (FGR; estimated fetal weight <10^th^ percentile) occurs in up to 10% of pregnancies in the developed world with suboptimal fetal nutrition and uteroplacental perfusion accounting for 25-30% of cases ^1, 2^. As one of the leading causes of stillbirth, miscarriage and infant morbidity, FGR is also associated with an increased risk of developing non-communicable diseases (NCDs; including cardiovascular disease, metabolic disease, central obesity, type 2 diabetes) in later life ^3-5^. Currently, there are limited predictive strategies to diagnose FGR, and there is no effective *in utero* treatment for FGR, thus no clinical intervention that could potentially impact or prevent the development of NCDs in adulthood.

The “developmental origins of health and disease” (DoHAD) hypothesis states that early environmental stressors during critical fetal developmental windows result in permanent, adaptive structural and physiologic changes that predispose the offspring to metabolic, endocrine, and cardiovascular disease in postnatal life ^6^. Animal models have shown that FGR is associated with increased hepatic gluconeogenic gene expression and glucose production in response to reduced placental glucose supply ^7-9^. Whilst this adaptive response may be beneficial temporarily *in utero*, persistent expression of these gluconeogenic genes beyond birth, when dietary glucose is no longer limited, may have adverse consequences to health with excess glucose production leading to hyperglycemia and the development of obesity and/or metabolic disease.

Liver gluconeogenesis is the *de novo* synthesis of glucose from non-carbohydrate precursors, and is principally controlled by activities of enzymes such as phosphoenolpyruvate carboxykinase (PCK/PEPCK) and glucose-6-phosphatase (G6PC) ^10^. There are two isozymes of PCK: the cytosolic isozyme PCK1 and the mitochondrial isozyme PCK2, both of which are involved in hepatic gluconeogenesis ^11^. Studies, predominantly performed in sheep models of FGR and focused on changes in late gestation, have shown fetal hepatic adaptations to support glucose production ^7, 12-14^. FGR is associated with increased expression of *Pck1, Pck2* and *G6pc* in the fetal liver, and transcriptomic analysis indicates increased amino acid catabolism and cell stress along with decreased mitochondrial activity ^13^. Moreover, increased expression of *Pck* has been shown to persist into adulthood, at the equivalent human age of 50-60 years ^15^. However, the effects FGR on liver gluconeogenesis around the initiation of FGR are lacking.

Given that fetal hepatic adaptations to glucose production are linked to insufficient placental glucose supply, directly targeting the dysfunctional placenta to increase glucose and other nutrient transport capacities may restore supply and mitigate DoHAD-associated developmental programming. We have developed the use of a polymer-based nanoparticle that facilitates non-viral, transient (does not integrate into the genome), gene delivery specifically to the placenta ^16, 17^. Using a biosynthetic HPMA-DMEAMA (N-(2-hydroxypropyl) methacrylamide-2-(dimethylamino)ethyl methacrylate) co-polymer, complexed with a plasmid containing the *human insulin-like 1 growth factor* (*hIGF-1*) gene under the control of trophoblast specific promotors (*PLAC1, CYP19A1*), we have successful shown efficient nanoparticle uptake into human syncytiotrophoblast *ex vivo* ^17^ as well as *in vivo* using animal models ^16, 18-20^. Short-term effects of *hIGF-1* gene therapy results in increased placental expression of glucose and amino acid transporters, maintenance of normal fetal growth under FGR conditions, and increased fetal glucose concentrations ^16, 18-20^. Importantly, our nanoparticle gene therapy is proven to be safe to both mother and fetus, and is capable of positively influencing placental function in diverse models of FGR as IGF-1 is central to most mechanisms responsible for FGR associated with placental dysfunction, and a major regulator of normal placental and fetal growth and development ^21^.

We have previously shown in the guinea pig MNR model of FGR, efficient uptake of our *hIGF-1* nanoparticle gene therapy into the guinea pig placenta, and no transfer of nanoparticle or *hIGF-1* plasmid into fetal circulation ^20^. In the present study, we aimed to characterize the effects of FGR on hepatic gluconeogenesis gene expression at the initial stages of FGR establishment in the fetal guinea pig, and determine whether treatment of the placenta with our *hIGF-1* nanoparticle gene therapy could resolve differences in hepatic gluconeogenesis gene expression in the FGR fetus.

## Materials and Methods

### Polymer synthesis and Nanoparticle formation

Detailed methods on the synthesis of the (PHPMA_115_-*b*-PDMAEMA_115_) copolymer, and nanoparticle formation can be found in *Wilson et al*., *2022* ^20^. Briefly, plasmids containing the *human IGF-1* gene under control of the trophoblast-specific *CYP19A1* promotor were mixed with the non-viral PHPMA_115_-*b*-PDMAEMA_115_ co-polymer for 1 h at room temperature to form the *hIGF-1* nanoparticle. Details about physiochemical properties, and cellular safety and efficiency of the PHPMA_115_-b-PDMEAMA_115_ nanoparticle has been previously published ^20^, and has been proven safe for both mother and fetus in numerous animal models ^16, 18, 19^. Finally, it is unlikely clinically that this nanoparticle gene therapy will be provided to a normally growing fetus. However, we have also shown that administration of this nanoparticle gene therapy to the placenta in a normal pregnancy environment results in down-regulation of decidual and placental mTOR signaling and growth factor gene expression in order to maintain placental homeostasis.

### Animal care and transuterine, intra-placental nanoparticle administration

Animal care and usage was approved by the Institutional Animal Care and Use Committees at Cincinnati Children’s Hospital and Medical Center (Protocol number 2017-0065) and University of Florida (Protocol number 202011236). Detailed information on animal care, maternal nutrient restriction (MNR) implementation, and ultrasound-guided transuterine, intra-placental nanoparticle administration can be found in *Wilson et al*., *2022* ^20^. Briefly, female (dams) Dunkin-Hartley guinea pigs (Charles River Laboratories, Wilmington MA) were purchased and assigned to either a control diet group (*n* = 7) where food and water was provided ad libitum, or MNR diet group (*n* = 12) where water was provided ad libitum, but food intake was restricted to 70% per kilogram body weight of the control group from at least four weeks preconception through to mid-pregnancy (GD30), thereafter increasing to 90%. Time mating with males was performed as outlined in *Wilson et al*., 2021 ^20^, and pregnancy confirmation ultrasounds performed at gestational day (GD) 21 using a Voluson I portable ultrasound machine (GE) with a 125 E 12 MHz vascular probe (GE). At GD30-33 dams underwent ultrasound-guided, transuterine, intra-placental injections of either *IGF-1* nanoparticle gene therapy (50 µg plasmid in 200 µL injection; *n* = 7 MNR diet) or sham injection (200 µL of PBS). Dams were sacrificed five days after injection (GD35-38) by carbon dioxide asphyxiation followed by cardiac puncture and exsanguination. Fetuses and placentas were removed from the gravid uterus and weighed. Glucose concentrations in both maternal and fetal blood was measured using a glucometer. Fetal sex was determined by examination of the gonads and confirmed using PCR as previously published ^19^. Fetal livers were dissected, weighed and halved to be either fixed in 4% w/v paraformaldehyde (PFA) or snap-frozen in liquid nitrogen and stored at -80ºC.

### Histology and Immunohistochemistry

For assessment of steatosis and fibrosis, 5 µm thick sections of PFA-fixed, paraffin embedded fetal liver tissue were obtained, de-waxed and rehydrated following standard protocol. Hematoxylin and eosin staining was performed as standard. Immunohistochemistry (IHC) was performed as previously described ^19^ to assess nuclear expression of Ki67 *(Invitrogen* MA5-14520; diluted 1:200), and nuclei were counter stained with hematoxylin. Following IHC staining, slides were imaged using the Axioscan (*Zeiss*) microscope, and 10 random 40x magnification images obtained using the Zen Blue software (*Zeiss*). For each image, cells positive for Ki67 (brown) and negative for Ki67 (blue) were counted using the Threshold and Watershed functions in ImageJ software ^22^, and averaged across the 10 images per slide to obtain a percentage positive result.

### RNA isolations and Quantitative PCR (qPCR)

For hepatic gene expression analysis, approximately 50 mg of snap frozen fetal liver tissue was lysed in RLT-lysis buffer (*Qiagen*) with homogenization aided by a tissue-lyzer. RNA was extracted using the RNeasy Mini kit (*Qiagen*), and included DNase treatment following standard manufacturers protocol. 1 µg of RNA was converted to cDNA using the High-capacity cDNA Reverse Transcription kit (*Applied Biosystems*) and diluted to 1:100. For qPCR, 2.5 µL of cDNA was mixed with 10 µL of PowerUp SYBR green (*Applied Biosystems*), 1.2 µL of KiCqStart SYBR Green Predesigned Primers (*Sigma*) at a concentration of 10 nM, and water to make up a total reaction volume of 20 µL. Gene expression was normalized using housekeeping genes *β-actin* and *Rsp20* (See ^20^). qPCR was performed using the Quant3 Real-Time PCR System (*Applied Biosystems*), and relative mRNA expression calculated using the comparative CT method with the Design and Analysis 2 v2.6.0 software (*Applied Biosystems*).

### Statistical Analysis

All statistical analyses were performed using SPSS Statistics 27 software with female and male fetuses analyzed separately. Generalized estimating equations were used to determine differences between diet and nanoparticle treatment. Dams were considered the subject, diet, and nanoparticle treatment treated as main effects, maternal environment treated as a random effect and gestational age as a covariate. Litter size was also included as a covariate but removed as there was no significant effect for any of the outcomes. Statistical significance was considered at P≤0.05. For statistically significant results, a Bonferroni post hoc analysis was performed. Results are reported as estimated marginal means ± standard error.

### Results

### Maternal nutrient restriction reduces fetal liver growth and results in brain-sparing at mid-pregnancy

The MNR diet resulted in decreased fetal weight in both females and males at mid-pregnancy, and there was no effect of *hIGF-1* nanoparticle treatment after 5 days (Fig. 1*a* & *b*). Fetal liver weight as a percentage of fetal weight was decreased in MNR and MNR + *hIGF-1* nanoparticle treatment in female and male fetuses compared to control (Fig. 1*c* & *d*). There was evidence of brain-sparing, as indicated by increased fetal brain:liver ratio in female and male fetuses, in the MNR and MNR + *hIGF-1* nanoparticle treatment fetuses when compared to control fetuses (Fig 1*e* & *f*).

**Figure 1.**
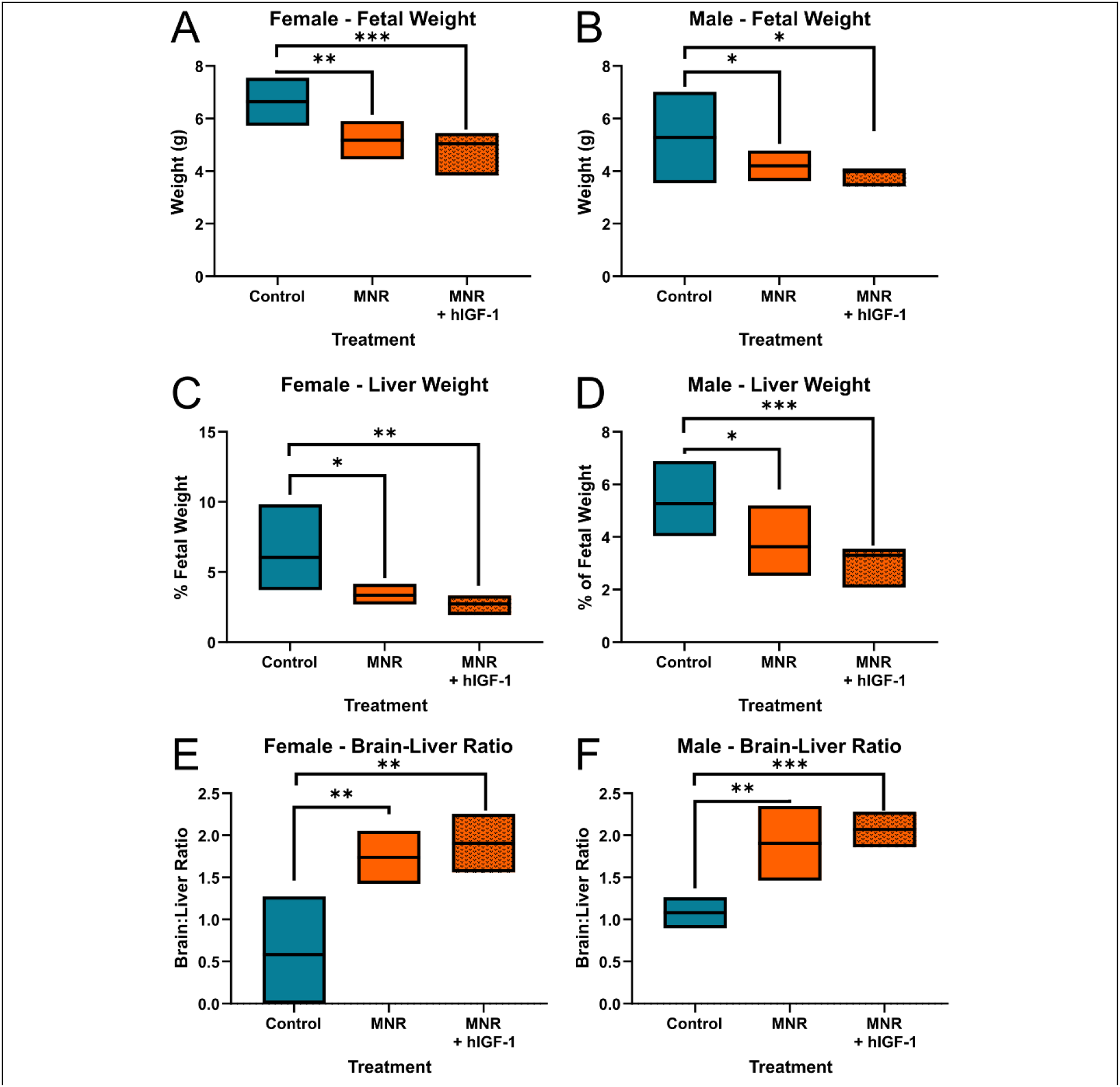
Effect of maternal nutrient restriction (MNR) and *hIGF-1* nanoparticle treatment on mid-pregnancy fetal growth parameters. **A**. MNR reduced mid-pregnancy fetal weight of female fetuses, and was not different with *hIGF-1* nanoparticle treatment. **B**. MNR reduced mid-pregnancy fetal weight of male fetuses, and was not different with *hIGF-1* nanoparticle treatment. **C**. MNR reduced mid-pregnancy fetal liver weight, as a percentage of fetal weight in female fetuses, and was not different with *hIGF-1* nanoparticle treatment. **D**. MNR reduced mid-pregnancy fetal liver weight, as a percentage of fetal weight in male fetuses, and was not different with *hIGF-1* nanoparticle treatment. **E**. MNR increased mid-pregnancy brain:liver weight ratio in female fetuses, and was not different with *hIGF-1* nanoparticle treatment. **F**. MNR increased mid-pregnancy brain:liver weight ratio in male fetuses, and was not different with *hIGF-1* nanoparticle treatment. *n* = 7 control dams (4 female & 8 male fetuses), 5 MNR dams (7 female & 7 male fetuses), and 7 MNR + *hIGF-1* nanoparticle dams (8 female & 11 male fetuses). Data are estimated marginal means ± 95% confidence interval. P values calculated using generalized estimating equations with Bonferroni post hoc analysis. *P<0.05; **P<0.01; ***P<0.001.

### Maternal nutrient restriction reduces liver proliferation in female fetuses, and is normalized with *hIGF-1* nanoparticle treatment

Morphologically, there was no evidence of increased steatosis or fibrosis in the livers of either female or male fetuses with MNR or *hIGF-1* nanoparticle treatment (Fig. 2). In female fetal livers, MNR reduced proliferation, as evident by the percentage of cells positive for Ki67, of the hepatocytes when compared to Control, however percentage of cells positive for Ki67 in MNR + *hIGF-1* nanoparticle treated was comparable to Controls (Fig. 3*a*). In male fetal livers, there was no effect of MNR or *hIGF-1* nanoparticle treatment on proliferation of hepatocytes (Fig. 3*b*).

**Figure 2.**
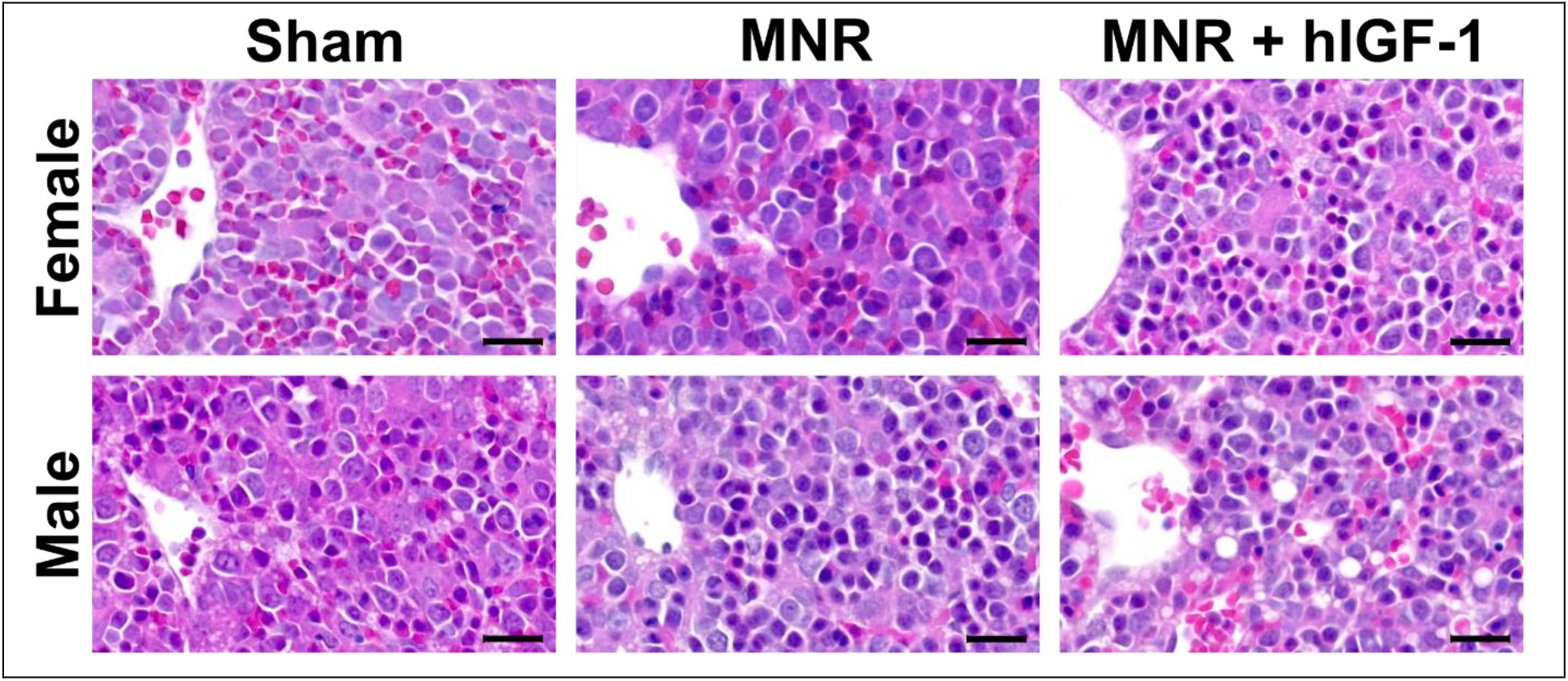
Representative images of hematoxylin and eosin (H & E) stained fetal livers at mid-pregnancy with maternal nutrient restriction (MNR) and *hIGF-1* nanoparticle treatment. There was no evidence of increased steatosis or fibrosis in the livers of either female or male fetuses with MNR or *hIGF-1* nanoparticle treatment. H & E images are taken at 40x magnification, scale bar = 10 µm. *n* = 4 control dams (4 female & 4 male fetuses), 5 MNR dams (5 female & 5 male fetuses), and 7 MNR + *hIGF-1* nanoparticle dams (7 female & 7 male fetuses).

**Figure 3.**
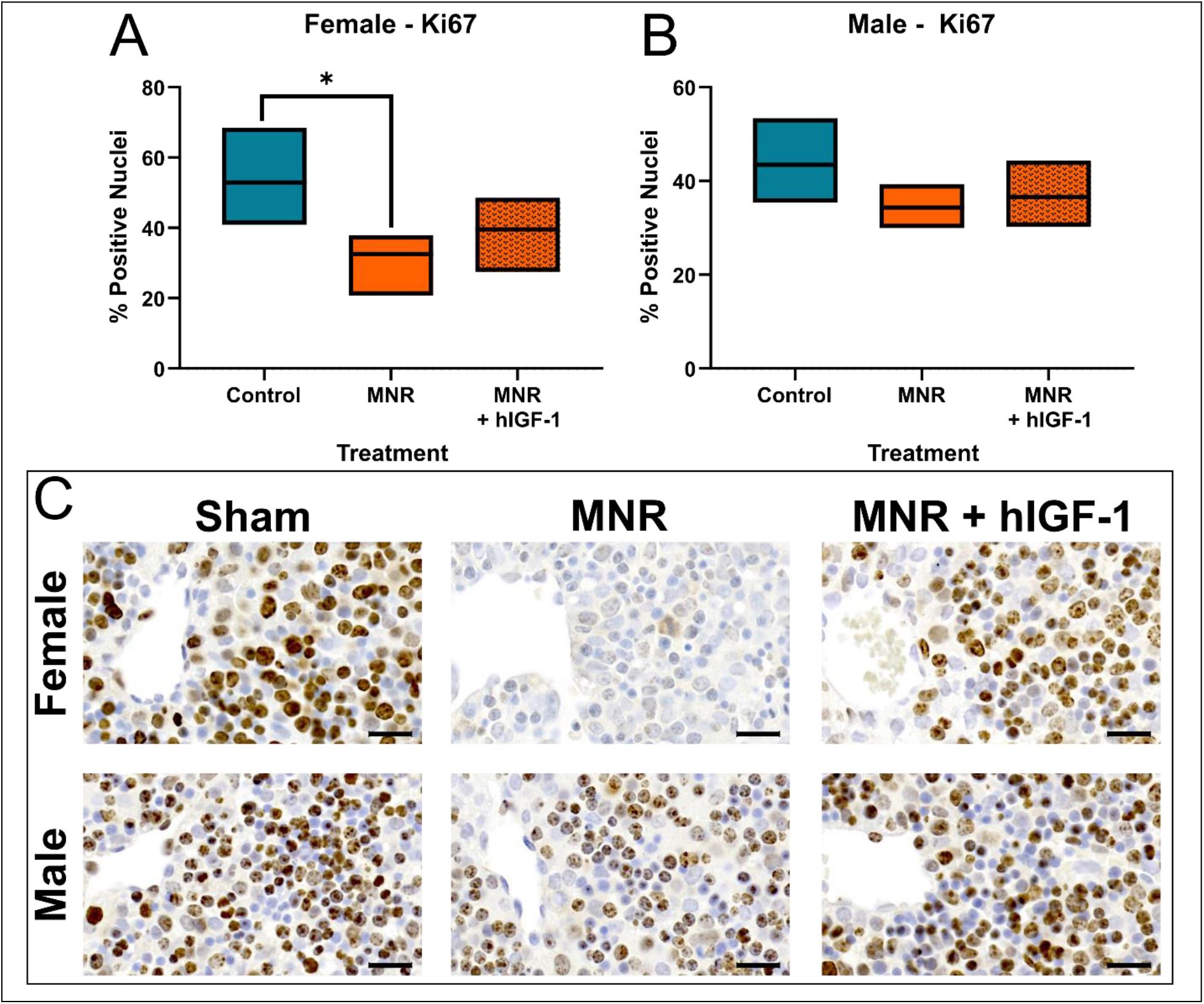
Effect of maternal nutrient restriction (MNR) and *hIGF-1* nanoparticle treatment on mid-pregnancy fetal liver hepatocyte proliferation. **A**. In female fetal livers, MNR reduced the percentage of hepatocytes positive for Ki67 (brown nuclei) when compared to Control. However, percentage of hepatocytes positive for Ki67 in MNR + *hIGF-1* nanoparticle treated female fetal livers was comparable to Control. **B**. There was no effect of either MNR or *hIGF-1* nanoparticle treatment on percentage of hepatocytes positive for Ki67 in male fetal livers. **C**. Representative images of Ki67 immunohistochemistry stained fetal livers, taken at 40x magnification; scale bar = 10 µm. *n* = 4 control dams (4 female & 4 male fetuses), 5 MNR dams (5 female & 5 male fetuses), and 7 MNR + *hIGF-1* nanoparticle dams (7 female & 7 male fetuses). Data are estimated marginal means ± 95% confidence interval. P values calculated using generalized estimating equations with Bonferroni post hoc analysis. *P<0.05.

### Expression of stress markers is reduced with *hIGF-1* nanoparticle treatment in MNR female fetal livers

In female fetal livers at mid-pregnancy, there was increased expression of growth factors *Tgfβ, Ctgf*, and *Mmp2* with MNR when compared to control (Fig. 4*a-c*). Expression of *Tgfβ, Ctgf*, and *Mmp2* remained increased in the MNR + *hIGF-1* nanoparticle treatment compared to control. On the other hand, expression of stress markers *Tnfα* and *Hif1α* were increased in MNR female fetal livers when compared to control, but decreased by *hIGF-1* nanoparticle treatment when compared to MNR, and towards comparable levels with control female fetal livers (Fig. 4*d* & *e*). qPCR analysis of growth factors and stress markers in the mid-pregnancy male fetal liver showed no differences in expression with either MNR or *hIGF-1* nanoparticle treatment (Supplemental Fig. 1).

**Figure 3.**
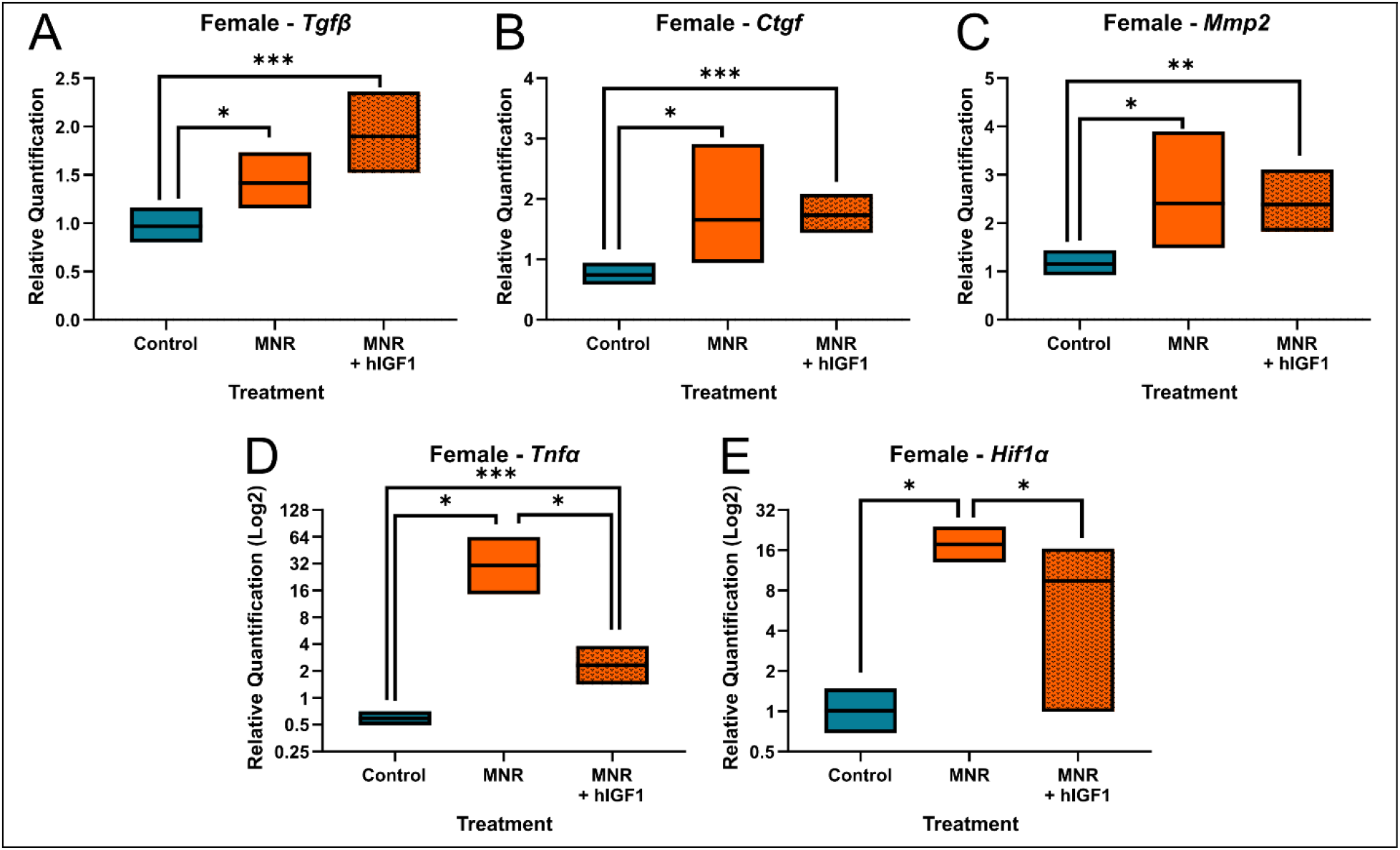
Effect of maternal nutrient restriction (MNR) and *hIGF-1* nanoparticle treatment on mid-pregnancy female fetal liver growth factor and stress marker gene expression. MNR increased expression of *transforming growth factor beta* (*Tgfβ*: **A**), *connective tissue growth factor* (*Ctgf*: **B**), *matrix metalloproteinase 2* (*Mmp2*; **C**) in female fetal livers compared to control female fetal livers. *hIGF-1* nanoparticle treatment did not affect expression of *Tgfβ, Ctgf*, and *Mmp2* which remained increased when compared to control. MNR increased expression of *tumor necrosis factor alpha* (*Tnfα*; **D**) and *hypoxia inducible factor 1 alpha* (*Hif1α*; **E**) when compared to control. *hIGF-1* nanoparticle treatment decreased expression in male MNR fetal liver tissue compared to MNR. *n* = 7 control dams (4 female fetuses), 5 MNR dams (7 female fetuses), and 7 MNR + *hIGF-1* nanoparticle dams (8 female fetuses). Data are estimated marginal means ± 95% confidence interval. P values calculated using generalized estimating equations with Bonferroni post hoc analysis. *P<0.05; **P<0.01; ***P<0.001.

### Placental *hIGF-1* nanoparticle treatment increased fetal glucose concentrations in male fetuses at mid-pregnancy

We have previously reported no difference in maternal blood glucose levels at mid-pregnancy with either diet or *hIGF-1* nanoparticle treatment ^20^. There was no difference in fetal blood glucose concentrations in females and males between MNR and sham (Fig. 5*a* & *b*). However, *hIGF-1* nanoparticle gene therapy increased fetal blood glucose concentrations when compared to control and MNR, but only in male fetuses.

**Figure 5.**
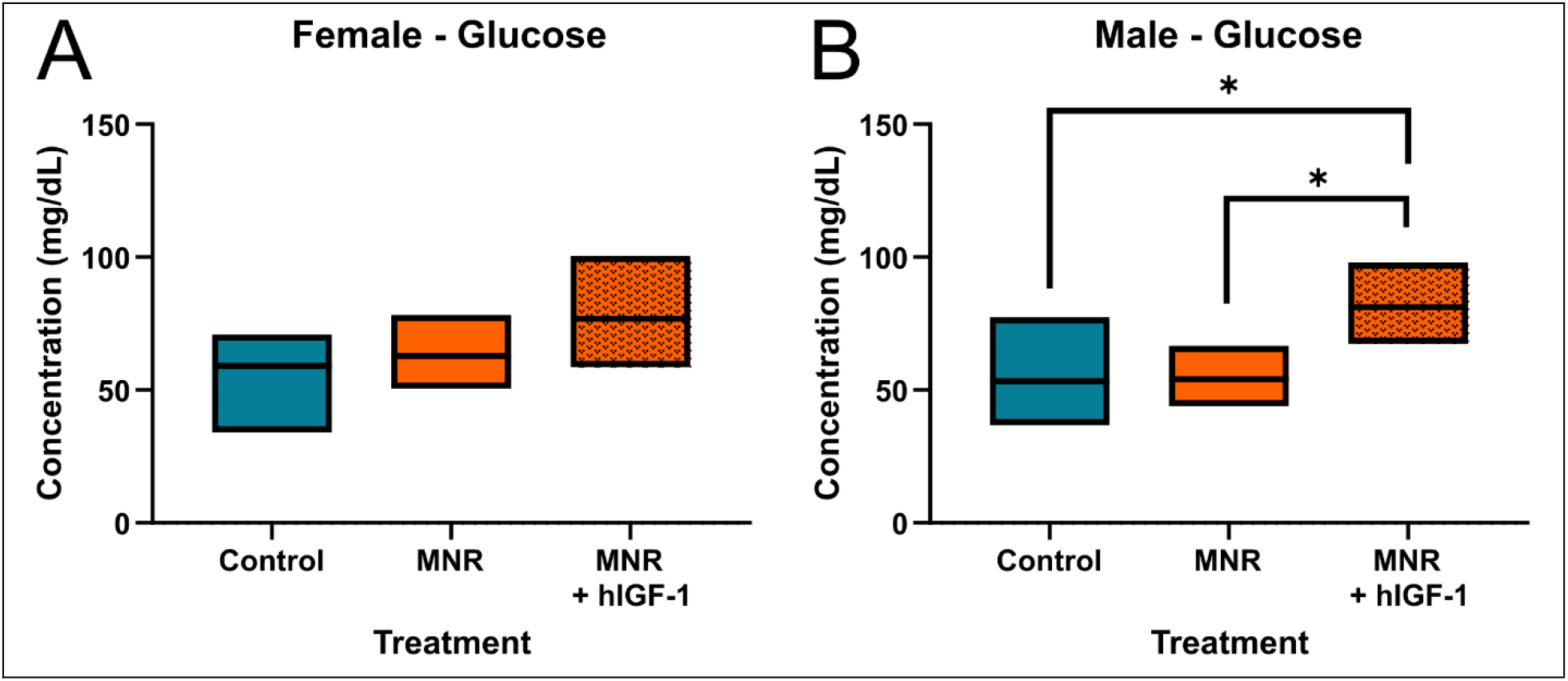
Effect of maternal nutrient restriction (MNR) and *hIGF-1* nanoparticle treatment on mid-pregnancy fetal blood glucose levels. **A**. There was no difference in blood glucose concentrations in female fetuses with either MNR or *hIGF-1* nanoparticle treatment. **B**. In male fetuses, there was no difference in blood glucose concentrations between control and MNR, but *hIGF-1* nanoparticle treatment increased blood glucose concentrations compared to sham. *n* = 7 control dams (4 female & 8 male fetuses), 5 MNR dams (7 female & 7 male fetuses), and 7 MNR + *hIGF-1* nanoparticle dams (8 female & 11 male fetuses). Data are estimated marginal means ± 95% confidence interval. P values calculated using generalized estimating equations with Bonferroni post hoc analysis. *P<0.05.

### Fetal liver gene expression of insulin-sensing and gluconeogenesis enzymes is affected by MNR at mid-pregnancy, and normalized with placental *hIGF-1* nanoparticle treatment in male fetuses only

It has been hypothesized that developmental programming in the fetal liver predisposing offspring to obesity and metabolic diseases, is due to increased gluconeogenesis in the fetal liver that persists after birth ^14, 23^. qPCR analysis of insulin-sensing and gluconeogenesis enzymes in the mid-pregnancy female fetal liver showed very few differences (Supplemental Fig. 2), with only *IgfBP3* increased in MNR compared to control, and returned to control expression levels in the MNR + *hIGF-1* nanoparticle group (Estimated marginal mean + SEM of Relative Expression: Control = 1.15 + 0.22 vs. MNR = 2.10 + 0.06 vs. MNR + *hIGF-1* nanoparticle = 1.31 + 0.22; P-value Diet = 0.002, P-value Treatment = 0.005). In mid-pregnancy males, MNR increased expression of *Igf-1*, and reduced expression of *Igf-2, G6pc* and *Pck1* when compared to control (Fig. 6*a-d*, respectively). Furthermore, MNR + *hIGF-1* nanoparticle treatment returned mid-pregnancy liver expression of *Igf-1, Igf-2, G6pc*, and *Pck1* to that, or towards that, of control. Additionally, *hIGF-1* nanoparticle treatment increased male fetal liver expression of *IgfBP1* and *GcgR* compared to control and MNR (Fig. 6*e* & *f*, respectively).

**Figure 6.**
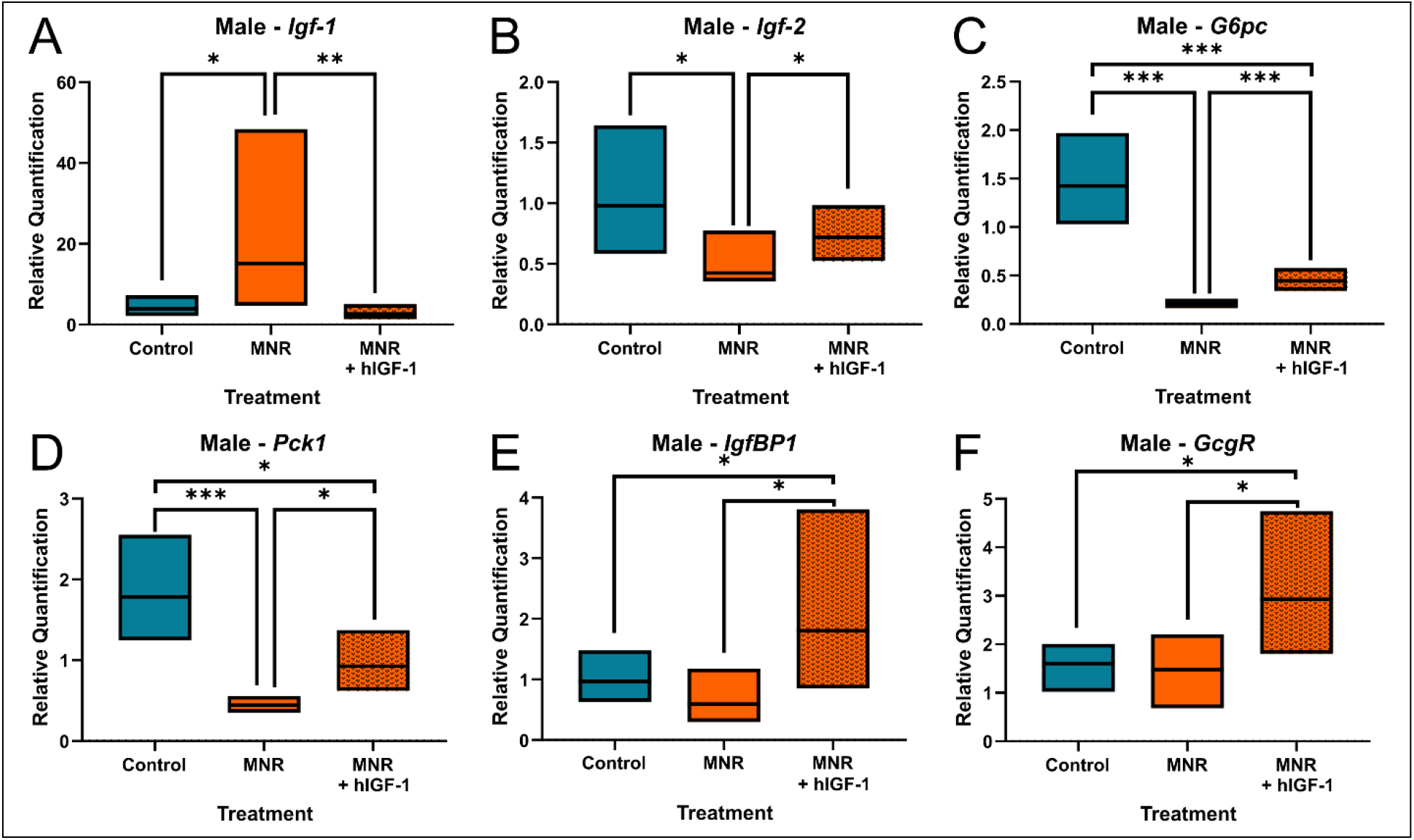
**Effect of maternal nutrient restriction (MNR) and *hIGF-1* nanoparticle treatment on mid-pregnancy male fetal liver insulin sensing and gluconeogenesis enzyme gene expression**. MNR increased expression of *insulin-like growth factor 1* (*Igf-1*: **A**), and decreased expression of *Igf-2* (**B**), *glucose-6-phosphatase* (*G6pc*; **C**), and *phosphoenolpyruvate carboxykinase 1* (*Pck1*; **D**) in male fetal livers compared to control male fetal livers. *hIGF-1* nanoparticle treatment restored expression of *Igf1, Igf2, G6pc* and *Pck1*, toward or back to normal. *hIGF-1* nanoparticle treatment increased expression of *Igf Binding Protein 1* (*IgfBP1*; **E**) and *Glucagon Receptor* (*GcgR*; **F**) compared to control and MNR male fetuses. *n* = 7 control dams (8 male fetuses), 5 MNR dams (7 male fetuses), and 7 MNR + *hIGF-1* nanoparticle dams (11 male fetuses). Data are estimated marginal means ± 95% confidence interval. P values calculated using generalized estimating equations with Bonferroni post hoc analysis. *P<0.05; **P<0.01; ***P<0.001.

## Discussion

Currently, there is no effective *in utero* treatment for FGR and thus, no clinical intervention that could potentially impact or prevent the increased risk of developing an NCD. In the present study, we show that MNR affects different physiological pathways depending on fetal sex; in females fetuses growth mechanisms are impacted compared to glucose production pathways in male fetuses. Moreover, we show short-term treatment of the placenta with *hIGF-1* nanoparticle gene therapy is capable of normalizing changes to fetal hepatic gene expression. Overall, this data provides further mechanistic understanding of how MNR and FGR affect hepatic gene expression and development and highlights the importance of understanding sex-specific risk windows during fetal development and potential considerations when developing pregnancy therapeutics.

The influence of fetal sex as a biological variable is well established. In human pregnancies, data consistently shows that risk of complications such as preterm delivery is higher in males compared to females ^24^. In this study, fetal weight, and liver weight were decreased in female and male fetuses with MNR when compared to control, an outcome routinely found in late-pregnancy in the guinea pig MNR model of FGR ^25-27^. However, at mid-pregnancy, and during the initial stages of FGR establishment, it was only in female fetal livers where MNR reduced liver hepatocyte proliferation, and increased the expression of growth factors and stress markers; expression remained comparable to Control in male livers. It has previously been reported in young adult male FGR offspring, that mRNA expression of *Tgfβ, Ctgf*, and *Mmp2* were increased in liver tissue ^28^, however, females were not assessed. Traditionally, increased expression of these growth factors has been associated with the promotion of liver fibrosis in adulthood ^29^, and increased expression would be considered detrimental to liver physiology. However, in mid-pregnancy fetal development, the pro-proliferative effects ^30^ are likely positive for organ development, particularly given reduced liver weight as a percentage of fetal weight. Overall, our results suggest in the early stages of FGR induction, growth pathways are more disrupted than gluconeogenic pathways in the livers of female fetuses, and that female fetuses prioritize supporting liver growth over glucose production.

Under normal growth conditions, endogenous fetal glucose production is absent because glucose supply from the placenta is sufficient. However, in cases of FGR the fetal liver increases gluconeogenesis and thus glucose production in order to maintain vital glucose supply to the developing organs ^14, 23^. Others have shown increased hepatic gene expression of gluconeogenesis enzymes in growth restricted fetuses in fetal sheep and rats in late-pregnancy, but did not separate fetal sex ^7, 8, 14^. In our study, at the time of FGR establishment, there was reduced expression of gluconeogenic enzymes *G6pc* and *Pck1*, which may represent a compensation trigger for the increased expression seen in late-pregnancy in other studies. However, the decrease in *G6pc* and *Pck1* expression was only observed in male fetuses. Adequate liver development and functionality ensures glycogen deposition and gluconeogenic ability, both of which are essential during the first stages of postnatal life ^31^. However, adaptations in liver development to adverse in utero environments, including blood flow and gene expression, whilst important to ensure short-term survival, may have longer-term detrimental consequences in the face of an enriched postnatal diet ^32^. Evidence from experimental models demonstrate responses to adverse in utero environments, with males affected to a greater extent than females, putting male offspring at higher risk of cardiovascular and metabolic disease ^33^. Overall, our data suggests that during the early stages of FGR establishment in males, insulin signaling and gluconeogenesis is affected disproportionately compared to growth patterning and may be a key contributor as to why males are at higher risk of developing obesity diabetes in adulthood.

At time of FGR establishment in both female and male fetuses, changes in liver gene expression of some genes were normalized with short-term placenta *hIGF-1* nanoparticle treatment. We have previously shown the inability for both nanoparticle and plasmid to cross the placenta and enter fetal circulation ^20^, thus any changes in fetal liver gene expression with placental *hIGF-1* nanoparticle treatment are indirect. During fetal development, there is multi-directional communication between mother, placenta and fetus ^34^. Nutrients, oxygen and signaling factors like hormones, are transferred across the placenta, through the umbilical cord to the liver. Thus, the liver is the first organ which receives nutrient and oxygen rich blood from the placenta ^35^. Presented here, there was increased expression of gluconeogenesis enzymes, towards normal, in male fetal livers with placenta *hIGF-1* treatment when compared to untreated. Analysis of the placental response to *hIGF-1* nanoparticle treatment shows increased expression of glucose and amino acid transporters ^20^, likely resulting in increased glucose transport across the placenta, presenting a potential mechanism by which fetal liver gene expression of these enzymes is changed. Furthermore, placental *hIGF-1* nanoparticle treatment resulted in reduced expression of hypoxia/stress markers in MNR female fetal livers. The molecular and physiological mechanisms behind the normalization of *Hif1α* and *Tnfα* are yet to be determined, however suggest the ability for the *hIGF-1* nanoparticle treatment of the placenta to result in reduced hypoxia in fetal livers. Our analysis of placental morphology with *hIGF-1* nanoparticle treatment indicates reduced interhaemal distance between maternal and fetal circulation likely resulting in increased oxygen diffusion ^20^, and a possible mechanism by which expression of *Hif1α* and *Tnfα* reduced.

The aim of this study was to assess the immediate, short-term impacts of placenta *hIGF-1* nanoparticle treatment on developmental programming in the fetal liver at the initial stages of FGR establishment. We have shown that secondary to placental treatment normalization of gene expression changes associated with MNR/FGR occurs. However, given the short time period, there was no significant improvement in fetal weight. Therefore, we are focusing our future research on performing multiple placental *hIGF-1* nanoparticle treatments over a longer time period in mid-late pregnancy. This data shows a potential method by which an *in utero* treatment of the placenta can impact developmental programming and may prevent increased risk of diseases like cardiovascular disease, metabolic disease, obesity and diabetes in later life.

## Acknowledgments

We would like to thank Drs Craig Duvall and Mukesh Gupta for providing the co-polymer, and Kristin Lampe for her assistance with the animal experiments.

## Contributions

RLW conceived the study, performed experiments, analyzed data and wrote manuscript. KKS performed experiments, analyzed data and edited manuscript. HNJ obtained funding, conceived the study and edited manuscript. All authors approve final version of manuscript.

## Ethics approval

Animal care and usage was approved by the Institutional Animal Care and Use Committees at Cincinnati Children’s Hospital and Medical Center (Protocol number 2017-0065).

## Data availability

All data needed to evaluate the conclusions in the paper are present in the paper and/or the Supplementary Materials.

