## Supplemental Material for "Placental nanoparticle gene therapy normalizes gene expression changes in the fetal liver associated with fetal growth restriction in a fetal sex-specific manner"

**Supplemental Data**

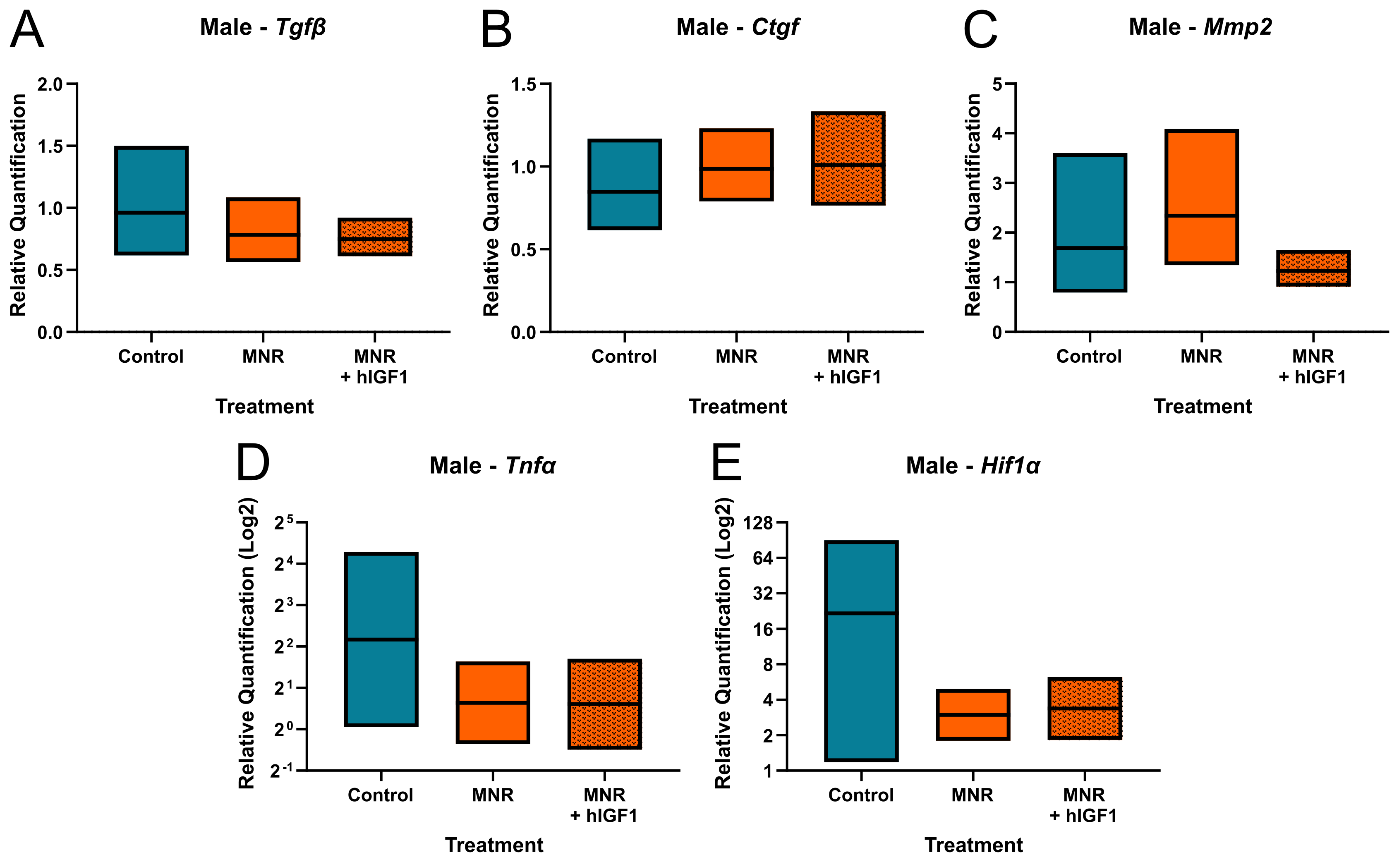

**Supplemental Figure 1. Effect of maternal nutrient restriction (MNR) and *hIGF-1* nanoparticle treatment on mid-pregnancy male fetal liver growth factor and stress marker gene expression.** There was no difference in the expression of *transforming growth factor beta* (*Tgfβ*: **A**), *connective tissue growth factor* (*Ctgf*: **B**), *matrix metalloproteinase 2* (*Mmp2*; **C**), *tumor necrosis factor alpha* (*Tnfα*; **D**) and *hypoxia inducible factor 1 alpha* (*Hif1α*; **E**) between Control, MNR and MNR + *hIGF-1*. *n* = 7 control dams (8 male fetuses), 5 MNR dams (7 male fetuses), and 7 MNR + *hIGF-1* nanoparticle dams (11 male fetuses). Data are estimated marginal means ± 95% confidence interval.

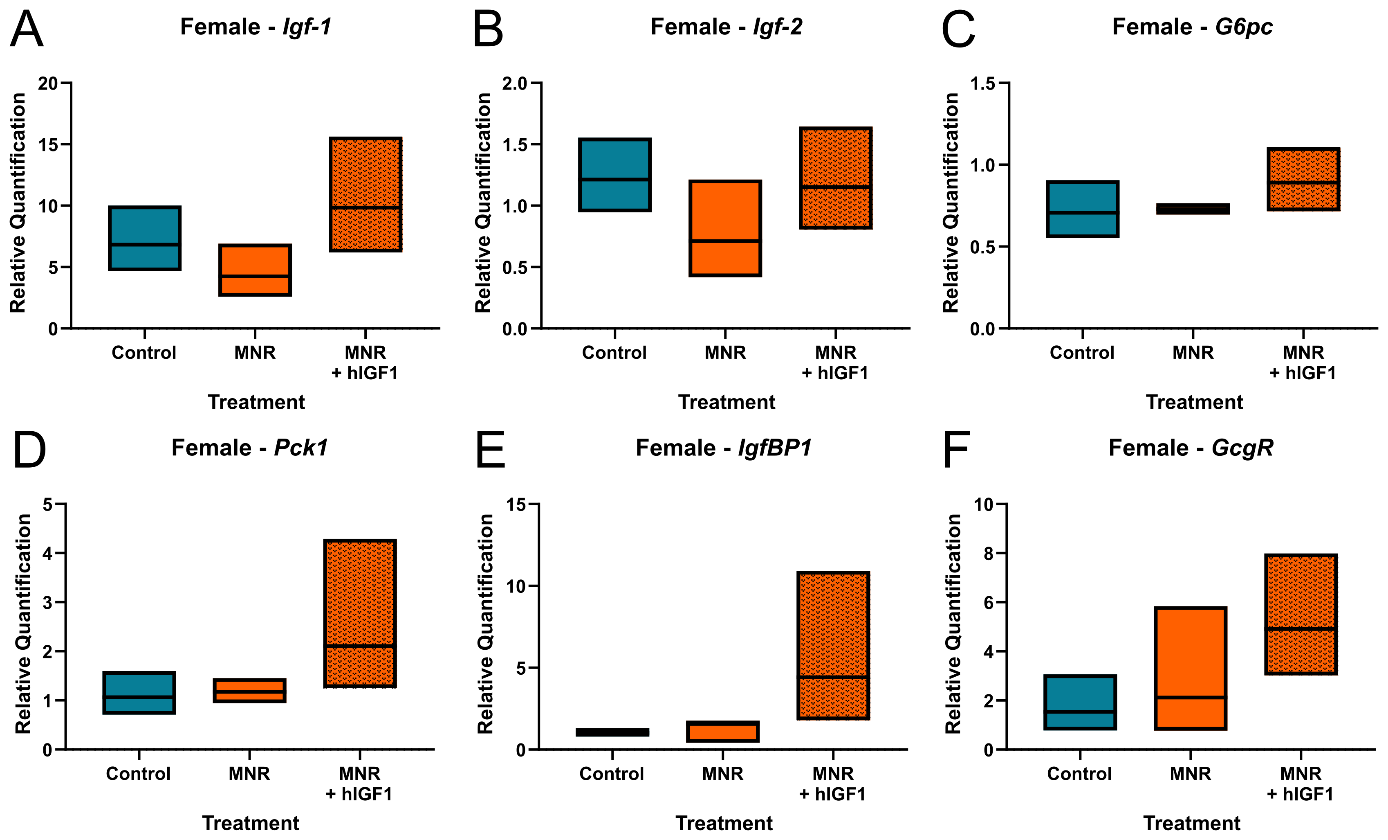

**Supplemental Figure 2. Effect of maternal nutrient restriction (MNR) and *hIGF-1* nanoparticle treatment on mid-pregnancy female fetal liver insulin sensing and gluconeogenesis enzyme gene expression.** There was no difference in the expression of *insulin-like growth factor 1* (*Igf-1*: **A**), *Igf2* (**B**), *glucose-6-phosphatase* (*G6pc*; **C**), *phosphoenolpyruvate carboxykinase 1* (*Pck1*; **D**) *Igf Binding Protein 1* (*IgfBP1*; **E**) and *Glucagon Receptor* (*GcgR*; **F**) between control, MNR and MNR + *hIGF-1*. *n* = 7 Control dams (4 female fetuses), 5 MNR dams (7 female fetuses), and 7 MNR + *hIGF-1* nanoparticle dams (8 female fetuses). Data are estimated marginal means ± 95% confidence interval.
